# Accurate detection of mosaic mutations at short tandem repeats from bulk sequencing data

**DOI:** 10.64898/2026.03.30.715227

**Authors:** Weixiang Wang, Wenlong Li, Chunyi Wang, Wenxuan Fan, Yonghe Xia, Xiaoxu Yang, Chong Chu, Yanmei Dou

## Abstract

Short tandem repeats (STRs) are among the most mutable regions of the human genome, yet their somatic mosaicism remains poorly characterized due to the technical challenges of distinguishing genuine mutations from high intrinsic polymorphism and sequencing noise. Here, we introduce BulkMonSTR, a computational framework that combines STR-specific error modelling with machine-learning classification to enable accurate detection of mosaic STR mutations from bulk next-generation sequencing data. BulkMonSTR identifies nucleotide-resolution mutations—including insertions, deletions, and single-nucleotide variants (SNVs)—and supports both control-independent and case-control study designs. Leveraging a comprehensive training dataset derived from pedigree-based validation and *in silico* spike-in simulations, our random forest classifier effectively discriminates true mosaic events from germline variants and technical artifacts. Benchmarking on simulated and real datasets demonstrates that BulkMonSTR achieves substantially improved precision and F1 scores across diverse coverages and variant allele frequencies. In normal samples, cancer samples and controlled *in silico* mixing experiments, BulkMonSTR consistently outperforms existing methods, capturing a broader spectrum of STR mutations—including those arising on non-reference alleles—while achieving high validation rates. By enabling systematic, genome-wide interrogation of STR mosaicism, BulkMonSTR provides a scalable foundation for investigating the contributions of somatic STR mutations to aging and disease.

## Introduction

STRs are repetitive elements consisting of tandem arrays of 1-6 base pair sequence motifs. These regions are highly mutable, primarily due to frequent DNA polymerase slippage during replication^1^. Based on pedigree analysis, the *de novo* mutation rate of STRs is approximately 1.1 × 10^−5^ *de novo* mutations per locus per generation—up to ~100-1,000-fold higher than that of SNVs^2^. STR mutations have been implicated in gene regulation and complex traits^3–7^ and are associated with a range of disorders, including more than 60 neurological conditions^8^ and tumorigenesis^9^.

Despite their functional significance, the landscape of mosaic STR mutations remains largely unexplored. Characterizing these mutations is technically challenging. Unlike somatic mutations in tumors, which often exhibit high variant allele frequencies (VAFs) due to clonal expansion, non-clonal mosaic mutations typically occur at very low VAFs. Identifying low-frequency mosaic STR mutations is complicated by multiple sources of technical noise, including PCR stutter errors^10^, sequencing errors^11^, and mapping errors. These challenges are further compounded by the lack of matched normal samples, which are critical for distinguishing true somatic mutations from germline variants.

Recent advances in small variant detection^12–15^, including image-based approaches leveraging deep learning models trained on comprehensive datasets and raw sequence alignment images, have achieved high sensitivity and precision in non-repetitive regions. However, these methods generally show modest sensitivity in STR regions, a limitation stemming from the polymorphic nature of STRs—where mutations may revert to the reference allele or shift between two non-reference alleles—event types that are overlooked by these methods. Moreover, these tools typically exclude multi-allelic and common population variants, further disregarding the highly mutable characteristics of STRs^16,17^. The recently developed prancSTR^18^ accounts for STR polymorphism and enables control-independent detection of mosaic STR mutations, but has the following limitations: (1) it does not leverage the rich information embedded in sequencing data for artifact identification; (2) it is limited to the detection of length changes.

Here we present BulkMonSTR, a computational framework for detecting mosaic STR mutations in bulk sequencing data. BulkMonSTR identifies nucleotide-resolution mosaic STR mutations, including insertions, deletions (indels), and SNVs, and is compatible with both control-independent and case-control study designs. By integrating diverse features tailored to STR regions, our framework effectively discriminates true mosaic mutations from germline variants and recurrent artifacts. Evaluations on simulated and real datasets demonstrate that BulkMonSTR achieves superior performance across diverse read coverages, variant allele frequencies, and sequencing technologies, providing a robust solution for accurately detecting mosaic STR mutations. BulkMonSTR is open source and available at https://github.com/douymLab/BayesMonSTR/tree/main/BayesMonSTR-BulkMonSTR.

## Result

### Overview of BulkMonSTR

BulkMonSTR executes three primary steps: identification of candidate STR alleles from sequencing data (Fig. 1A), likelihood-based genotyping (Fig. 1B), and machine learning (ML)-based mutation classification (Fig. 1C).

**Figure 1.**
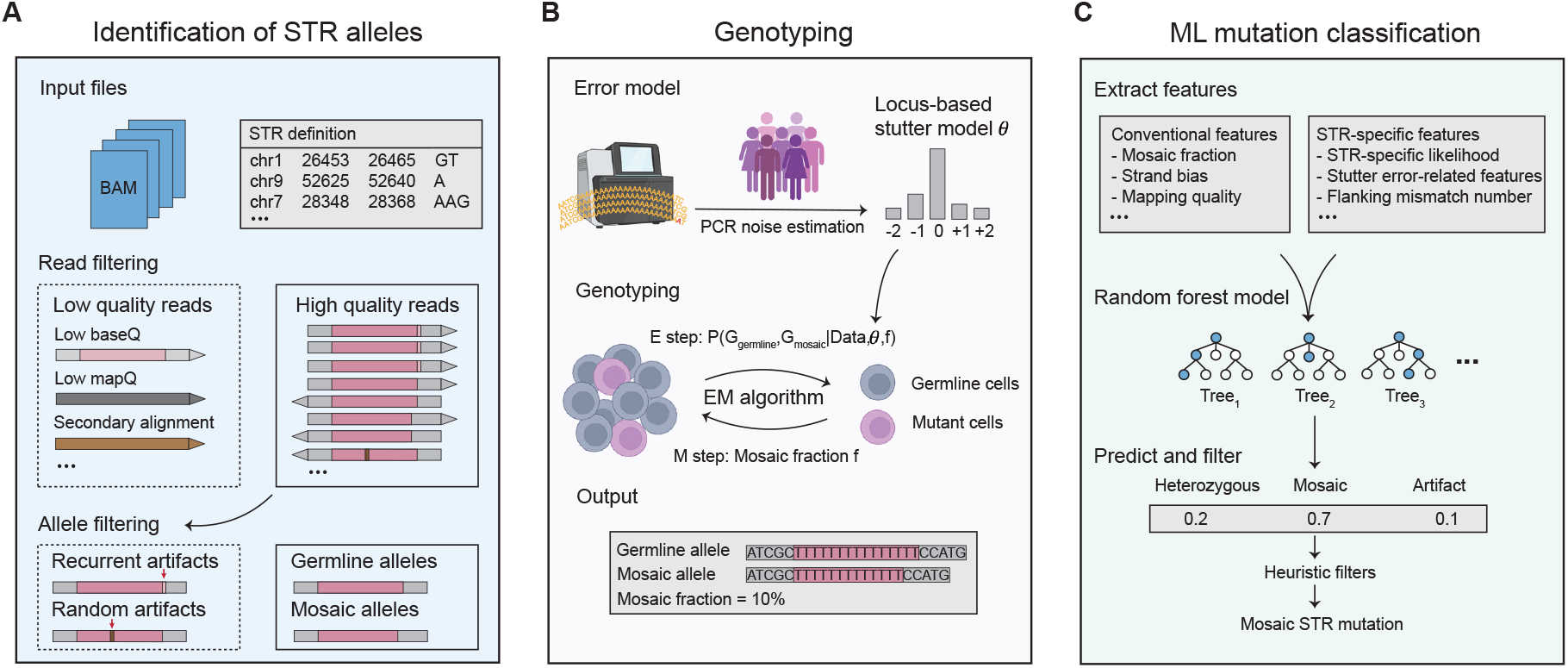
Overview of the BulkMonSTR workflow. (**A**) STR allele identification module. BulkMonSTR takes aligned BAM files and predefined STR regions as input. Reads spanning target STR loci are extracted and subjected to read-level and allele-level filtering to generate high-quality candidate alleles. (**B**) The framework of probabilistic genotype inference. A locus-specific stutter error model is estimated from population data and incorporated into the genotyping framework. An EM algorithm is then applied to estimate mutant allele fraction and infer maximum-likelihood genotypes. (**C**) Machine learning-based mutation classification. Candidate mosaic mutations are filtered using a random forest model trained based on a rigorously curated training set, followed by heuristic filters to further improve precision.

To identify STR alleles, BulkMonSTR first extracts candidate alleles by segmenting spanning reads aligned to target STR from a widely used STR panel containing ~1.6 million STR loci^19^, retaining the repeat sequence together with a 5-bp window of flanking sequence on each side as the initial candidate allele. A major challenge in STR allele identification is the presence of recurrent, non-independent technical noise. For example, mismatch errors that are prevalent in STR regions often mask true mutation signals (Supplementary Fig. 1A-C). To mitigate this issue, we implemented a two-step filtering strategy consisting of read-level and allele-level filtering to generate high-confidence candidate alleles. Read-level filtering removes reads that fail predefined quality-control criteria, such as low mapping quality or secondary alignment (Methods), thereby reducing potential alignment and sequencing artifacts. Allele-level filtering further suppresses recurrent mismatch artifacts, based on empirical observations from real sequencing data where such artifacts exhibit lower base quality at mismatched bases and strong strand bias (Methods, Supplementary Fig. 1D-F). This strategy improved genotyping sensitivity while reducing false positive rates and computational runtime in simulated datasets (Supplementary Fig. 1G-I).

For genotype inference, BulkMonSTR applies an Expectation-Maximization (EM) algorithm, making genotype inference from sequencing reads based on the high-confidence candidate alleles identified in the previous step (Fig. 1B, Methods). To account for technical noise, an STR-specific stutter error profile was first estimated across all analyzed samples (Methods). This profile was then used to quantify locus-specific background error rates and was incorporated directly into the probabilistic EM framework.

Finally, to further ensure high-fidelity mutation calling, BulkMonSTR employs a random forest classification model, leveraging diverse read-level features to distinguish true mosaic mutations from artifacts and germline variants. To this end, BulkMonSTR incorporates both conventional features—such as VAF, strand bias, and mapping quality—and STR-specific features, such as STR-region stutter error patterns and STR-flanking mismatch counts (Supplementary Table S1). The classifier was trained on a curated dataset spanning a range of sequencing depths and VAFs. It categorizes candidate mutations as mosaic mutations, artifacts, or germline heterozygous variants. Furthermore, additional hard filters—such as population allele frequencies from the STR reference panel^20,21^ (EnsembleTR Version II, 3,550 individuals)—are applied to further eliminate residual germline variants (Methods).

### Machine learning based mutation classification

To construct a comprehensive training dataset, we employed two complementary approaches: (1) pedigree-based mutation classification and (2) *in-silico* spike-in simulations of mosaic STR mutations (Methods, Fig. 2A, Supplementary Fig. 2). For the pedigree-based approach, we used an Ashkenazi Jewish trio (HG002: son, HG003: father, HG004: mother) from the Genome In a Bottle (GIAB) project^22^, with both Illumina sequencing data and Element sequencing data for each person (Supplementary Table S2). Candidate mutations were first identified by the BulkMonSTR genotyping module and prancSTR, and then parental sequencing data were used to distinguish inherited germline variants from *de novo* mutations. Furthermore, orthogonal cross-platform sequencing data were used to identify artifacts (Methods). This strategy identified 589,383 artifacts, 260,360 germline heterozygous variants, and 775 mosaic STR mutations (representative examples of HG002 mosaic STR mutations are shown in Supplementary Fig. 3) across different read coverages (Fig. 2A). For the spike-in approach, mosaic mutations were simulated with BamSurgeon^23^ (v1.4.1) across a wide range of sequencing depths, VAFs, and indel lengths (Fig. 2B-C). In total, 321,781 simulated mosaic STR mutations (comprising 206,896 Indels and 114,885 SNVs) were generated (Fig. 2A, Methods, Supplementary Table S3). Together, these datasets formed a training set integrating real and simulated mutations, comprising 1,172,299 sites in total.

**Figure 2.**
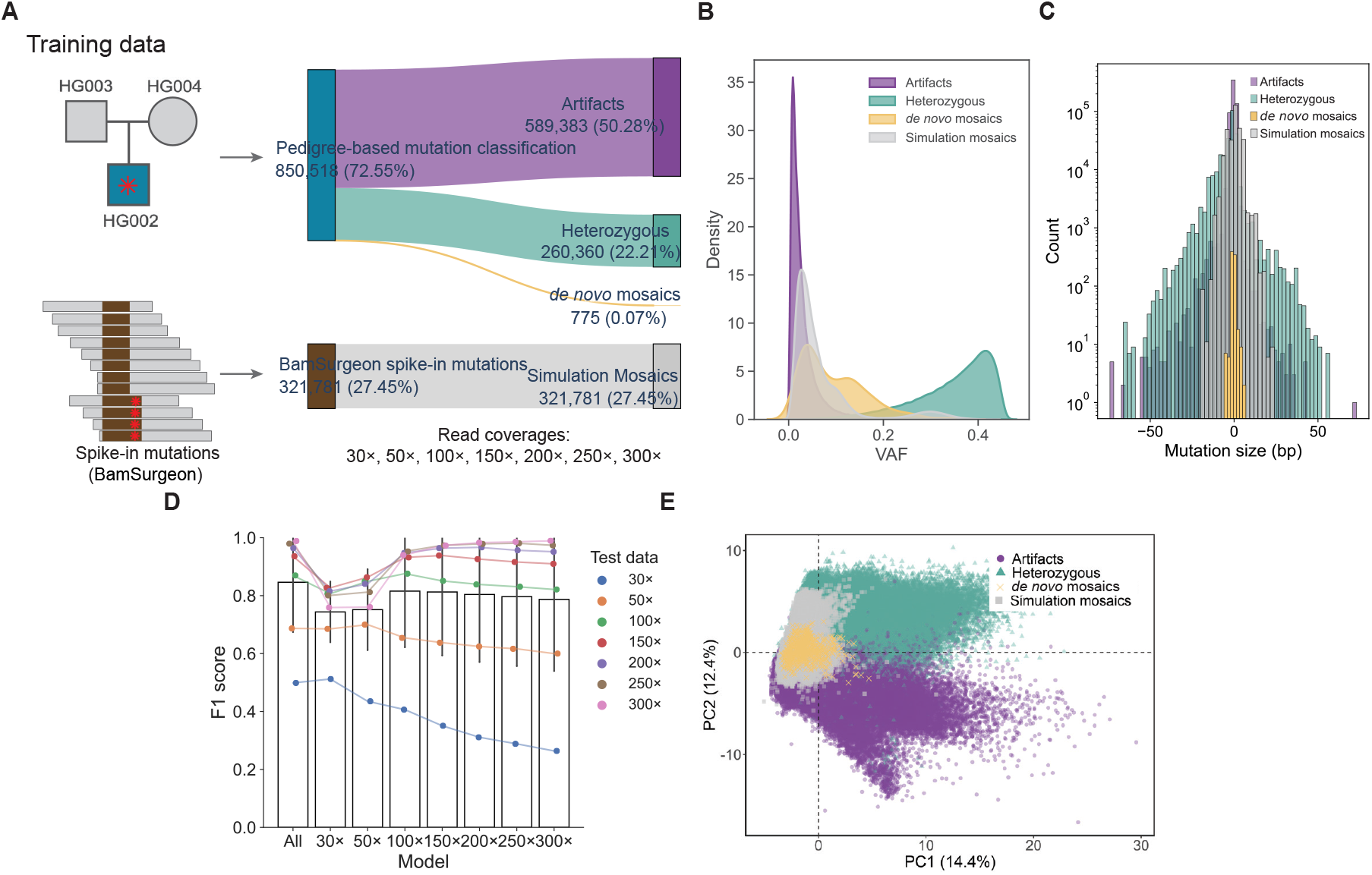
Construction of an RF Model Using a Rigorous Training Set. (**A**) Schematic overview of training dataset construction via pedigree-based mutation classification and spike-in simulations (see details in Methods). Cross-validated and simulated mutation types are color-coded as follows: artifacts (purple), heterozygous (green), mosaics (yellow), and simulated mosaics (grey), with the same scheme applied in **B, C, E**. (**B**) VAF distributions of different-category mutations. Compared with mosaic mutations, artifacts are of lower VAFs, and heterozygous variants have much higher VAFs. (**C**) Mutation size distributions. (**D**) The RF model trained on reads of varying coverages in general achieves comparable or higher F1 scores than models trained on single-coverage data. (**E**) PCA based on read-level features shows that spike-in simulated mosaic mutations and pedigree-based *de novo* mosaic mutations form a single distinct cluster, clearly separated from technical artifacts and germline heterozygous mutations. Each point represents a locus projected into two-dimensional PCA space.

Using this high-quality training dataset, we trained seven random forest (RF) models on data from different read coverage levels (ranging from 30× to 300×), as well as a consolidated model trained on pooled data from all coverages. The consolidated model achieved consistently higher or comparable F1 scores on independent test sets across all coverage levels and was therefore selected for all subsequent benchmarking analyses (Fig. 2D). Analysis of feature importance in the selected model revealed that mosaic fraction, STR-specific genotyping likelihood, flanking mismatch number, and stutter-related features were the strongest contributors to classification performance (Supplementary Fig. 4).

To visually assess whether our simulation strategy produced realistic mosaics and to evaluate the discriminative power of the read-level features, we performed a principal component analysis (PCA). This analysis was performed using the same feature set as that used in our RF model. In the resulting PCA space, the spike-in simulated mosaic mutations and pedigree-based *de novo* mosaic mutations formed a single, distinct cluster. This cluster was clearly separated from both technical artifacts and germline heterozygous mutations (Fig. 2E). The tight clustering of the two independent mosaic sets validates that our *in silico* spike-in strategy effectively recapitulates genuine mosaicism, while their clean separation from other variant classes visually confirms the strong discriminative power of the used read-level features. Of note, our pedigree-based approach substantially expands the catalog of mosaic variants in the GIAB HG002 sample, increasing it by approximately two-fold—from 85^24^ to 238 sites (Supplementary Fig. 5, Supplementary Table S4).

### Benchmarking mosaic STR detection using real bulk sequencing data

We evaluated BulkMonSTR on two Illumina whole-genome sequencing (WGS) datasets: high-coverage data from the GIAB HG005 sample (300×) and moderate-coverage data from 170 TCGA blood samples^25^ (35-55×). For HG005, we performed cross-validation using both trio sequencing data and orthogonal Element sequencing data from the same GIAB reference material (Methods, Supplementary Table S2). For the TCGA blood samples, we employed read-based haplotype phasing^15,26,27^ for mutation cross-validation, which is a standard approach for identifying false-positive mutation calls in somatic mosaicism research (Methods).

In both datasets, BulkMonSTR achieved significantly higher precisions (HG005, *P* = 1.14 ×10^−110^; TCGA blood samples, *P* = 6.92×10^−13^; one-sided Fisher’s exact test) and F1 scores (HG005, *P* = 2.23×10^−308^; TCGA blood samples, *P* = 1.1×10^−4^; bootstrap test) than prancSTR (Fig. 3A, Methods). Expanded pedigree-based analysis revealed that over 15% of mutations identified by prancSTR in HG005 were inherited germline variants, whereas BulkMonSTR efficiently excluded these, with only ~3% of its calls validated as inherited germline variants (Fig. 3B-C). Consistently, phasing analysis of TCGA samples demonstrated that BulkMonSTR more effectively excluded heterozygous variants phased as ‘Hap = 2’ compared to prancSTR, with a substantially lower Hap = 2 fraction (33% vs. 66%) (Fig. 3D–E; Supplementary Fig. 6). Beyond germline variants, we found that the BulkMonSTR RF model successfully excluded 78% of the artifactual mutations detected by prancSTR (Supplementary Fig. 7A). Expanded analysis using SHAP^28^ revealed that mosaic fraction, STR-specific likelihood, flanking mismatch level, and stutter-related features (e.g., the difference between mutant VAF and the estimated stutter error rate) were key factors of BulkMonSTR in excluding these false-positive loci (Supplementary Fig. 7B).

**Figure 3.**
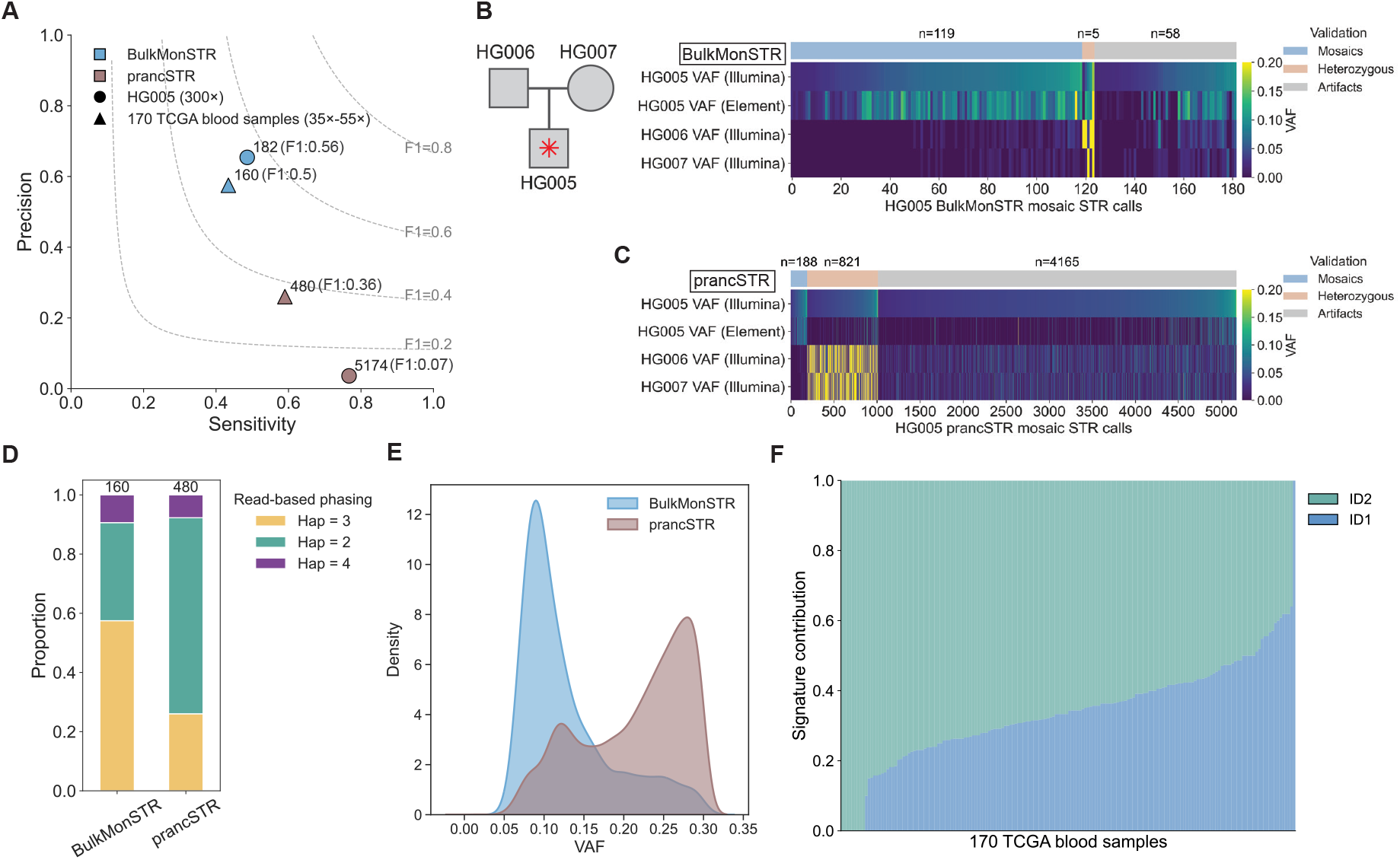
Benchmarking mosaic STR detection using real bulk sequencing data. (**A**) BulkMonSTR consistently achieves higher F1 scores and precisions than prancSTR in high-coverage (300×) HG005 WGS data and 170 moderate-coverage (35×-55×) TCGA blood samples. Blue and brown indicate BulkMonSTR and prancSTR, respectively; circles denote HG005 and triangles denote TCGA blood samples. (**B**-**C**) VAFs of candidate mosaic STR mutations detected by BulkMonSTR (**B**) and prancSTR (**C**) in parental sequencing data and orthogonal Element sequencing data. HG006 and HG007 correspond to the father and mother of HG005, respectively. The color bar indicates the magnitude of VAFs. Validation results are indicated above the heatmaps. (**D**) Read-based phasing in 170 TCGA blood samples shows that BulkMonSTR achieves a higher validation rate than prancSTR, with a greater proportion of loci phased as “Hap = 3” (58% vs. 26%). (**E**) The VAF distributions of mosaic STR calls identified by BulkMonSTR and prancSTR in 170 TCGA blood samples. PrancSTR detects a higher proportion of candidates with VAF > 0.2, consistent with its increased fraction of calls phased as “Hap = 2”. (**F**) Mutational signatures of mosaic STR mutations (indels) across 170 TCGA blood samples are dominated by COSMIC ID1 and ID2 related to replication slippage processes.

We further investigated the indel mutational signatures across 170 TCGA blood samples. The mutation spectrum was predominantly attributed to COSMIC signatures ID1 and ID2^29,30^ (Methods), which are associated with replication slippage at repetitive sequences. This finding aligns with the STR-specific mutational mechanism and further supports the validity of the BulkMonSTR mutation calls (Fig. 3F).

Element sequencing data has been shown to exhibit significantly lower sequencing error rates in repetitive genomic regions than Illumina, offering a distinct advantage for high-resolution analysis of STR regions^31–33^. To evaluate whether the current model of BulkMonSTR, trained on Illumina sequencing data, is suitable for mosaic STR detection in Element sequencing data, we analyzed the precision of mutation calling in HG005 and demonstrated that the current model trained with Illumina achieved a high validation rate of 75% (Supplementary Fig. 8, Methods).

### Benchmarking mosaic STR detection using in-silico mixed bulk sequencing data

To further evaluate BulkMonSTR’s performance in mutation detection across varying read depths and VAFs, we performed *in silico* read mixing using a tumor sample (HG008) from GIAB^34^. The HG008 sample consists of DNA from a high-purity pancreatic ductal adenocarcinoma (PDAC) tumor cell line and its matched normal cells. To assess performance across sequencing depths and variant allele fractions, we constructed synthetic mixtures by combining tumor and normal BAM files at defined ratios to simulate varying tumor purities, followed by down-sampling to a range of coverages (Supplementary Fig. 9A, Methods). BulkMonSTR was then run on these mixtures in both control-independent and case-control modes.

We first benchmarked BulkMonSTR against prancSTR in control-independent mode. Compared to prancSTR, BulkMonSTR demonstrated significantly higher precision (*P* = 4.4 × 10^−11^, one-sided Wilcoxon signed-rank test) across a wide range of sequencing depths and tumor purities (Fig. 4A, Supplementary Fig. 10), along with greater accuracy in estimating simulated tumor purities (Supplementary Fig. 9B). Specifically, BulkMonSTR consistently achieved > 70% precisions at tumor purities of 5%-60% (VAF 0.025-0.3, Methods). In addition, BulkMonSTR also demonstrated significantly higher sensitivity (*P* = 4.4 × 10^−11^, one-sided Wilcoxon signed-rank test) than prancSTR (Fig. 4A, Supplementary Fig.11). Across all simulated depths and purities, BulkMonSTR detected 19,372 mosaic STR mutations, 17,056 (88%) of which were validated as true positives. In contrast, prancSTR identified 6,143 mutations, with 4,911 (80%) validated (Fig. 4B). Overall, BulkMonSTR achieved a significantly higher F1 score than prancSTR (*P* = 1.5 × 10^−11^, one-sided Wilcoxon signed-rank test; Fig. 4C).

**Figure 4.**
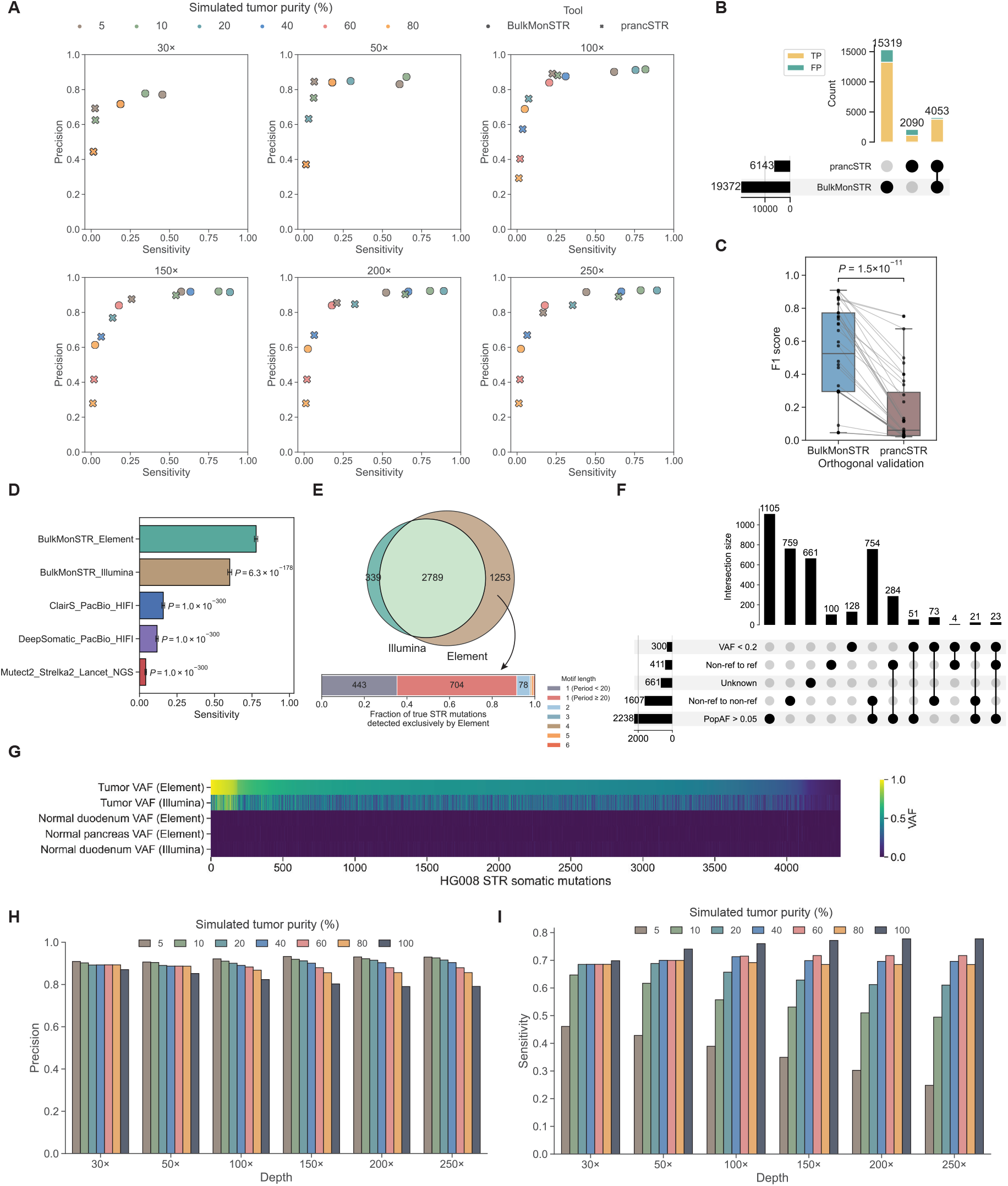
Benchmarking mosaic STR detection using *in-silico* mixed bulk sequencing data. (**A**) BulkMonSTR (control-independent mode) achieves higher sensitivity and precision than prancSTR across a range of sequencing depths and simulated tumor purities. (**B**) BulkMonSTR (control-independent mode) detects more mosaic STR mutations and shows a higher validation rate than prancSTR. Orthogonal validation results are color-coded: true positives in yellow and false positives in green. (**C**) BulkMonSTR (control-independent mode) achieves a significantly higher F1 score than prancSTR across sequencing depths and simulated tumor purities. Gray solid lines connect F1 scores at matched tumor purities and sequencing depths. Statistical comparison between BulkMonSTR and prancSTR was performed using a one-sided Wilcoxon signed-rank test. (**D**) BulkMonSTR (case-control mode) shows approximately 5-fold higher sensitivity than other small-variant callers. Error bars represent 95% confidence intervals estimated using binomial sampling. P-values were calculated using a binomial test. (**E**) BulkMonSTR (case-control mode) detected more true-positive mosaic STR mutations in Element sequencing data (n = 4,042 loci) than in Illumina data (n = 3,128 loci). Loci uniquely detected in Element sequencing data are more frequently located in homopolymer regions of ≥ 20 bp, consistent with the lower error rate of Element sequencing in these regions. (**F**) STR mutations uniquely detected by BulkMonSTR (case–control mode) highlight its ability to accommodate diverse allelic backgrounds, thereby contributing to increased sensitivity. (**G**) STR mutations (n = 4,381) detected by BulkMonSTR (case-control mode) show high reliability, with VAFs close to 0.5 in tumor samples (PDAC) and near zero in normal samples (normal duodenum and pancreas). (**H**-**I**) BulkMonSTR (case-control mode) shows robust precision (**H**) and sensitivity (**I**) across varying sequencing depths and simulated tumor purities.

We next benchmarked BulkMonSTR (case-control mode) against Mutect2^16^, Strelka2^17^, and Lancet^35^—three state-of-the-art methods for calling somatic mutations from NGS data—as well as ClairS^14^ and DeepSomatic^12^, which are based on long-read technologies. Detecting somatic mutations from the original tumor-normal pair, BulkMonSTR exhibited substantial overlap with these callers: it shared 180 mosaic STR mutations calls exclusively with DeepSomatic, 65 loci with both DeepSomatic and ClairS, and 51 loci with all tools (Supplementary Fig. 9C). Notably, compared with these well-established mutation callers, BulkMonSTR demonstrated markedly higher sensitivity, reaching 77.8% in Element data and 60.2% in Illumina data (Fig. 4D)—approximately five-fold higher than other tools in STR regions—while maintaining high validation rates (77% in Element and 83% in Illumina sequencing data, Supplementary Fig. 9D). Furthermore, while we observed a high concordance rate between validated mosaic STR mutations identified by BulkMonSTR from Illumina and Element sequencing data (Fig. 4E), BulkMonSTR detected ~30% more such mutations from Element data, likely due to improved detection of mutations occurring at long (≥20 bp) homopolymer loci (Fig. 4E).

Further analysis of the enhanced sensitivity of BulkMonSTR in mutation detection shows that relative to other callers, BulkMonSTR identified 3,963 unique validated mosaic STR mutations across Illumina and Element data. A substantial fraction of these calls reflected the polymorphic nature of repeat regions: mutant alleles at 56% sites had a population allele frequency >5% (Methods); 41% sites involved mutations from one non-reference allele to another (representative examples shown in Supplementary Fig. 12); and 10% sites are mutations from a non-reference allele to the reference allele (Fig. 4F). Importantly, the majority of these somatic STR mutations detected by BulkMonSTR showed VAFs of approximately 0.5 in tumor samples (Illumina and Element) and near-zero VAFs in matched normal tissues across platforms, supporting their high confidence (Fig. 4G).

Notably, the variant call sets obtained from the GIAB FTP repository^36^—generated by Mutect2 ^16^, Strelka2^17^, and Lancet^35^—did not include SNVs and indels with a ≥1% population allele frequency. This filtering strategy, however, is ill-suited for STR regions, where high allelic diversity is a fundamental property, and likely compromises detection sensitivity. Furthermore, current deep learning–based approaches such as DeepSomatic^12^ and ClairS^14^ typically only incorporate channels representing differences from the reference genome in their training framework, which likely limits their sensitivity for detecting mutations occurring on non-reference alleles.

Finally, we evaluated performance across varying tumor purities and sequencing depths. In the case-control mode, precision consistently exceeded 0.77 across sequencing depths and tumor purities (Fig. 4H). Sensitivity was robust across sequencing depths, maintaining ≥50% sensitivity at tumor purities ≥5% (Fig. 4I). Together, these results demonstrate the robust performance of BulkMonSTR in case-control mode across diverse experimental settings.

### BulkMonSTR captures the full spectrum of STR mutations

To evaluate BulkMonSTR’s ability to comprehensively characterize STR mutations, we further analyzed the mutation spectra of the GIAB reference samples (HG002, HG005, HG008) and 170 TCGA blood samples. Unlike conventional tools that restrict mutation detection to the human reference allele^12,16^, our method models the full allele pool at each locus. This approach is essential for highly polymorphic, hypermutative STRs, allowing us to capture mutations on diverse allelic backgrounds. Indeed, we found that at least 15% of indel mutations and a substantial fraction of mismatches within STRs occurred on non-reference alleles, reflecting the true allelic diversity at these loci (Fig. 5A-C). For instance, we identified a 6-bp deletion on a non-reference allele in a TCGA blood sample (Fig. 5D), as well as a complex case where an A>T mismatch arose on a non-reference allele already bearing a 1-bp deletion and an SNV (Fig. 5E).

**Figure 5.**
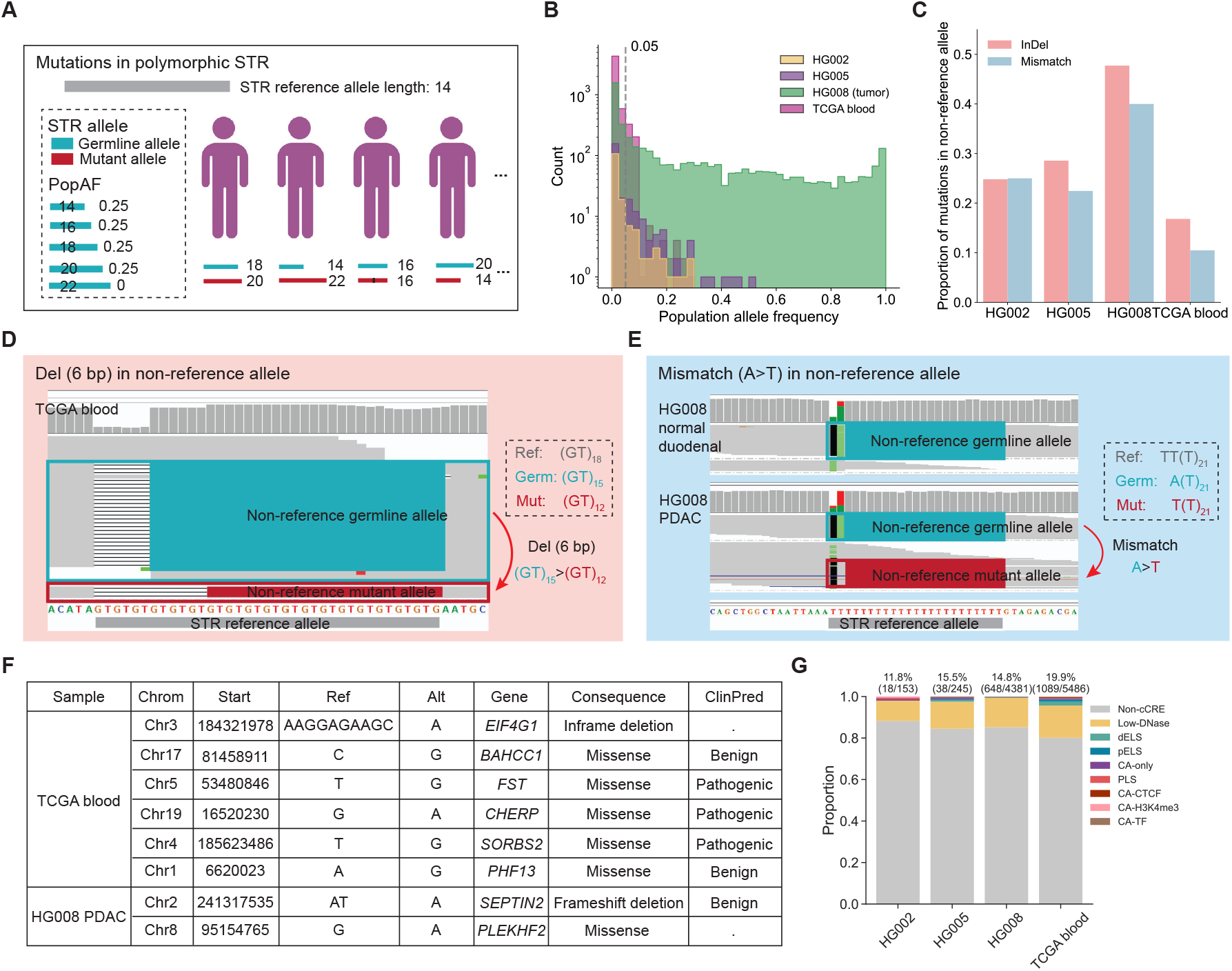
BulkMonSTR captures the full spectrum of STR mutations. (**A**) A schematic illustrating that STR mutations frequently arise on diverse allelic backgrounds due to high polymorphism and mutability. (**B**) Cross-validated STR mutations have diverse population allele frequencies (population allele frequencies from EnsembleTR calls on 3,550 individuals from 1000 Genomes Project and H3Africa^20,21^; Methods). (**C**) Cross-validated STR mutations are of diverse allelic backgrounds. (**D-E**) IGV^[34]^ plots showing a 6-bp deletion (**D**) in a non-reference allele in a TCGA blood sample and an A>T mismatch (**E**) in a non-reference allele in the HG008 PDAC sample. Colors indicate reads supporting different alleles: green for the non-reference germline allele, red for the non-reference mutant allele, and gray for the STR reference allele. The dashed box contains the sequences of the reference, germline, and mutant alleles at this locus. (**F**) Summary of STR mutations in protein-coding regions. (**G**) Functional annotation of STR mutations shows that >10% of STR mutations are located in putative candidate cis-regulatory element (cCRE) regions (See Methods in detail). Numbers above the bars indicate the proportion and number of STR mutations in cCRE regions.

A key advantage of BulkMonSTR is its nucleotide-resolution detection, which enables simultaneous identification of both indels and mismatches—a significant improvement over length-based methods. This resolution also mitigates alignment artifacts; in the complex example above, conventional methods would likely misinterpret the mutation as a germline variant due to misalignment of non-spanning reads. By treating each STR locus as an independent haplotype unit and considering only fully spanning reads, BulkMonSTR avoids such artifacts, enabling sensitive and accurate mutation detection.

The functional relevance of this enhanced resolution is evident in downstream analyses. While length-based approaches like prancSTR miss mismatches entirely, BulkMonSTR identified eight STR mutations in coding regions, six of which were mismatches. Four of these mismatches were predicted to be pathogenic, underscoring the critical importance of nucleotide-resolution for assessing functional impact (Fig. 5F). For noncoding STR mutations, we leveraged ENCODE4 cCRE annotations and found that 11.8–19.9% of these variants overlapped candidate cis-regulatory elements, pointing to their potential regulatory roles (Fig. 5G).

Together, these results demonstrate that BulkMonSTR captures the full spectrum of STR mutations—from diverse allelic backgrounds to single-nucleotide changes—providing a comprehensive view of mutational complexity and functional consequences that is unattainable with conventional or length-based methods.

## Discussion

We have developed BulkMonSTR, a method that enables accurate, nucleotide-resolution detection of STR mutations from bulk sequencing data. By addressing key limitations of conventional tools—namely, their restriction to reference alleles or reliance on length-based analysis—BulkMonSTR provides a comprehensive view of STR mutational diversity.

A key advantage of BulkMonSTR is its high accuracy, achieved through STR-specific error modeling and a machine learning framework trained on a robust dataset. Across benchmarks, our method achieves 55-80% precision in detecting mosaic mutations, several-fold higher than existing approaches while maintaining competitive sensitivity. This accuracy enables confident downstream analyses linking mosaic STR mutations to health and disease.

BulkMonSTR also offers nucleotide-resolution detection, a critical advance over length-based methods such as prancSTR. By capturing both indels and mismatches, our method enables a more complete characterization of STR mutation spectra and mechanisms, particularly for mosaic mutations. Additionally, by modeling the full allele pool at each locus rather than restricting to reference alleles, BulkMonSTR embraces the hypermutative, polymorphic nature of STRs, allowing identification of diverse mutations on non-reference backgrounds.

We acknowledge current limitations. The model is trained on Illumina NGS data with a limited training set; expanding to additional sample types and sequencing platforms—such as long-read technologies—will further improve generalizability and enable characterization of STRs in regions challenging for short reads.

Together, these capabilities position BulkMonSTR as a powerful tool for robust and comprehensive STR mutation detection in human health and disease research.

## Methods

### Identification of candidate alleles

The first step of BulkMonSTR involves extracting candidate STR alleles from sequencing alignment files. Given the frequent mismatch errors in repeat regions, which lead to inconsistent indel presentations across reads even after BWA^37^ left-alignment, BulkMonSTR treats each STR as a unified haplotype rather than relying on alignment-position-based calling. STR alleles are extracted by trimming all spanning reads aligned to the target STR, including the repeat sequences and 5 base pairs of flanking sequences. To minimize genotyping errors caused by noise alleles, we employ three progressive noise reduction strategies: (1) Low-quality read filtering: Reads that meet any of the following criteria are filtered out: (i) secondary alignments, (ii) supplementary alignments, (iii) improper paired alignments, (iv) mapping quality (MapQ) lower than 20, (v) average base quality lower than 20, (vi) mismatch fraction greater than 0.1, (vii) marked as duplicates, or (viii) flagged as QC-fail. (2) Single-read allele exclusion: To eliminate random noise, any candidate allele supported by only a single read is excluded. (3) Recurrent mismatch artifact filtration: To address pervasive mismatch artifacts from sequencing (e.g., phasing noise^11^), we designed a dedicated algorithm. This algorithm assumes that a less-supported allele is likely an artifact derived from a more-supported allele of the same length with the lowest edit distance (defined as the number of mismatches between the two alleles). The algorithm includes two main components: (i) Artifact derivation identification: For each candidate allele, we identify its most probable artifact source allele based on the criteria mentioned above. (ii) Artifact confirmation criteria: An allele is classified as an artifact if it meets both of the following conditions: (a) The average base quality of its mismatch bases is less than 20, or the proportion of mismatch bases with quality less than 20 exceeds 50%; (b) The allele is supported exclusively by reads from a single strand. Alleles confirmed as artifacts based on these criteria are subsequently filtered out.

### Estimation of the stutter error model

A core principle of the BulkMonSTR algorithm is to determine whether the observed fraction of a mosaic allele is significantly higher than the stutter error rate. Accurate estimation of the stutter error model for each STR locus is therefore essential. We assume that the simultaneous occurrence of mosaic mutations at the same STR locus across multiple individuals is exceedingly rare. With a sufficient number of samples, this enables precise estimation of the per-locus stutter error model, minimizing stutter error over-estimation and preserving sensitivity to true mosaic events. Similar to the approach in HipSTR^19^, we employ a generalized stepwise error model to characterize per-locus stutter errors. As the sample genotypes are initially unknown, they are treated as hidden variables. An EM algorithm is then used to iteratively estimate the stutter model parameters. The EM algorithm continues until the maximum number of iterations (t = 500) is reached, or the parameter change falls below 0.0001, yielding a stutter error model for each STR locus. Given that the STR length of the true allele is *a*, the probability of observing a read with STR length *r* is:

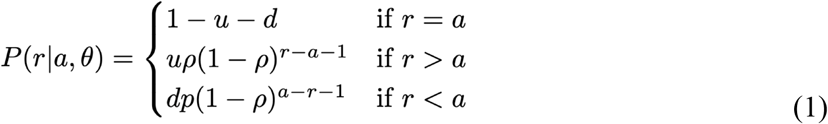

The stutter model *θ* comprises three parameters: the probability of stutter adding (*u*) or removing (*d*) repeats from the true allele in an observed read, and a geometric distribution with a parameter *ρ* that governs the size of the stutter-induced changes.

### Mosaic genotyping

After extracting candidate alleles, we compute the most likely mosaic genotypes. We assume that independent mutations are unlikely to recur at the same STR locus. For each STR locus, we hypothesize the presence of two cell types in the bulk sequencing sample: germline cells, which carry genotypes inherited from the parents, and mutant cells, which have undergone a mutation and carry genotypes distinct from the germline source. The proportion of mutant cells is defined as the mosaic fraction (*f*), an unknown parameter that depends on the occurrence timing of the mutation and the process of clonal selection. Therefore, the mosaic genotyping process is modeled as the estimation of the mosaic fraction, given the unknown mosaic genotypes. Once the mosaic fraction is estimated, the maximum likelihood mosaic genotypes are derived as the final result. This represents a parameter estimation problem involving latent variables, making it well-suited for the EM algorithm. The EM algorithm is employed to iteratively estimate the mosaic fraction and solve for the maximum likelihood mosaic genotypes.

Since the EM algorithm is sensitive to initial values, we test 20 initial mosaic fraction values, ranging from 0 to 1 in increments of 0.05, to prevent convergence to a local optimum. The initial value that maximizes the likelihood of the mosaic genotypes is selected. The EM algorithm then iterates through the following steps:

The E-step computes the mosaic genotype likelihood using the following formula:

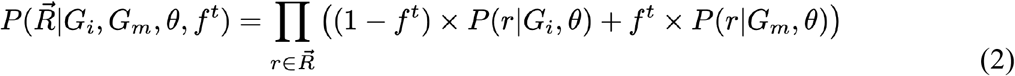

Where:

- *G*_*i*_ is the individual’s germline genotype.
- *G*_*m*_ is the genotype of the mutant cell.
- 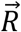 represents all sequencing reads.
- *θ* is the previously estimated locus-based stutter model, which includes both an in-frame stutter error model (*u*_*in*_, *d*_*in*_, *p*_*in*_) for stutters that are integer multiples of the motif length and an out-of-frame stutter error model (*u*_*out*_, *d*_*out*_, *p*_*out*_) for non-integer multiples of the motif length.
- *f*^*t*^ is the mosaic fraction at the t-th iteration.
- *r* represent a given sequencing read.

The likelihood of a read *r* originating from a cell with a specific genotype is given by:

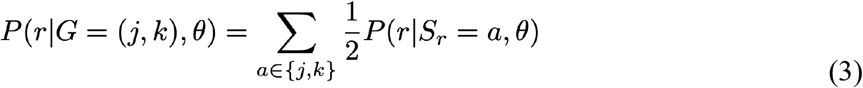

Where:

- *j,k* represent two given STR haplotypes.
- *S*_*r*_ represents the alleles or cells from which the read *r* originates.
- Other parameters are defined in Equation (2).

The likelihood of read *r* originating from a specific allele *a* is given by:

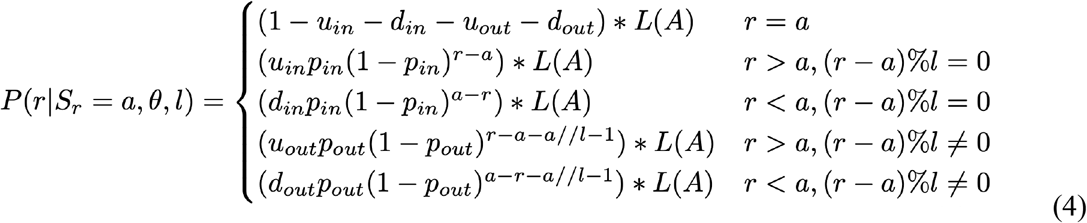

Stutter errors include indels that are integer multiples of the motif length *l*, as well as those that are not. We distinguish between these two types of stutter errors by modeling them separately as an in-frame stutter error model with parameters *u*_*in*_, *d*_*in*_, *p*_*in*_ and an out-of-frame stutter error model with parameters *u*_*out*_, *d*_*out*_, *p*_*out*_. The STR-specific alignment likelihood *L*(*A*) assumes that a read differs from the underlying haplotype by at most one indel, as in previous studies^19,38^. Given a haplotype *X* of length *L*_*X*_, and assuming that read *Y* with length *L*_*Y*_ is free of stutter errors, the alignment likelihood *L*(*A*) is computed based on the agreement between each base *i* in the read and its corresponding base in the haplotype:

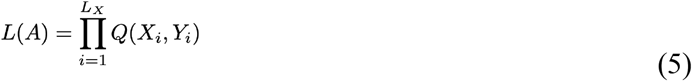

Alternatively, if read *Y* contains a stutter deletion relative to the haplotype *X* with deletion size Δ*L*, we assume that the deletion can occur at any position *k* within the haplotype, where *k* ∈ 1, …, *L* − Δ*L* + 1. Given a deletion starting at position *k*, the alignment likelihood *L*(*A, k, ins*) is defined as:

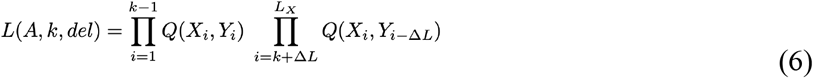

Finally, if read *Y* is assumed to result from a stutter insertion relative to haplotype *X*, with insertion size Δ*L*. We assume that the inserted sequence arises from a local duplication of adjacent bases. The insertion is allowed to occur at any position *k* within the haplotype, where *k* ∈ 1, …, *L* + 1. Given an insertion occurring at position *k*, the alignment likelihood *L*(*A, k, del*) is defined as:

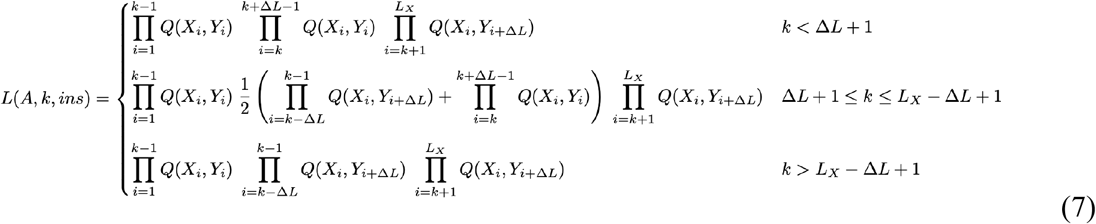

For reads assumed to contain a stutter insertion or deletion event, the overall alignment likelihood is computed as the arithmetic mean of the likelihoods across all possible indel positions:

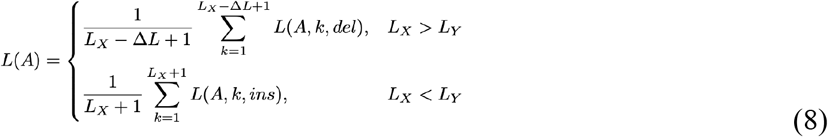

*Q*(*X*_*i*_, *Y*_*i*_) denotes the probability of observing read base *Y*_*i*_ given the underlying haplotype base *X*_*i*_:

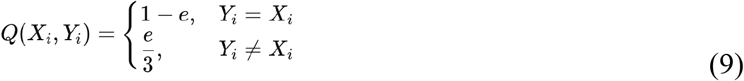

Where:

- *e* indicates sequencing error rate.

The M-step is then used to update the mosaic fraction based on these probabilities:

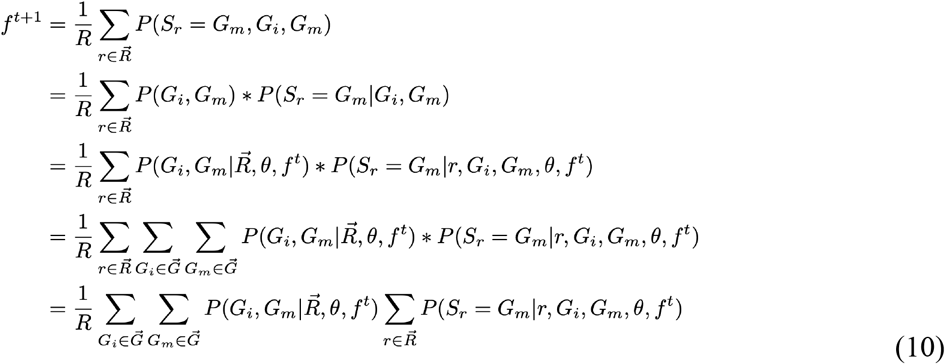

Where:

- *R* represents the number of all sequencing reads.
- Other parameters are defined in Equation (2-3).

The posterior probability that the read *r* originates from mutant cells is computed using the following formula:

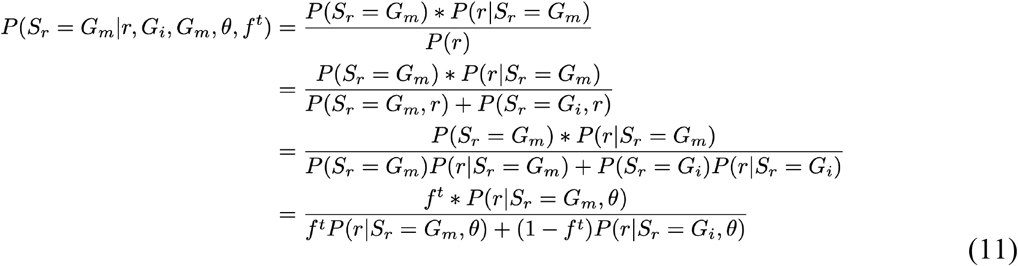

Intuitively, the update rule for the mosaic fraction involves computing the fraction of times a read originates from a mutant cell. The EM algorithm is iterated until either the maximum number of iterations (t = 500) is reached or the change in parameters is less than 0.0001. Finally, based on the estimated mosaic fraction, the maximum likelihood sequence-based mosaic genotypes are derived.

### Extraction of read-level features

Following mosaic genotyping with BulkMonSTR, both germline and mosaic alleles are identified, and the corresponding mosaic genotypes are determined. All raw reads from the BAM file are subsequently extracted. Given the presence of stutter errors, the alleles observed in the reads may not match the determined mosaic genotypes. To address this, each read is assigned to its most probable originating allele by calculating the STR-specific likelihood as Equation 4. A total of 51 features were extracted for indel model training and prediction, while 60 features were used for mismatch model training and prediction. These features are classified into the following categories: likelihood-related, VAF-related, stutter-related, mapping-related, baseQ-related, mismatch-related, and other features. A detailed description of these features is provided in Supplementary Table S1. The genotype likelihoods under the homozygous, heterozygous, and mosaic are formulated as follows:

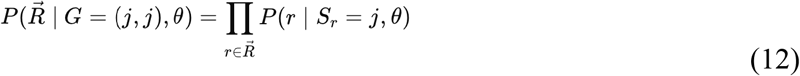

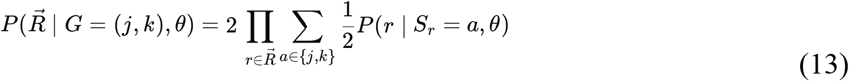

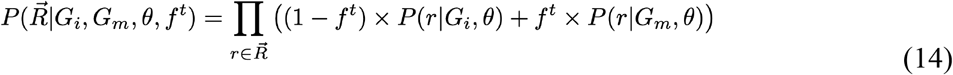

### Training of the random forest model

The training datasets included mosaic mutations, germline heterozygous, and artifacts from HG002, and simulated mosaic mutations from SimData1 (Supplementary Fig. 2). These loci were down-sampled to various depths, including 30×, 50×, 100×, 150×, 200×, 250×, and 300×. Features were extracted based on the results of BulkMonSTR mosaic genotyping, and these features were subsequently used for model training and testing. The dataset was split into a 9:1 ratio for training/validation and testing, respectively. The training/validation sets underwent 10-fold cross-validation with a grid search to optimize the hyperparameters. The model was then refitted using the optimized hyperparameters on the entire training/validation dataset, and its final performance was evaluated on the independent test set. Separate models were trained for each coverage, as well as a consolidated model trained across all depths. These models were tested on independent test data from different depths. Ultimately, the consolidated model, trained on all depths, was selected for subsequent benchmarking.

### Pedigree-based mutation classification

We labeled mosaic, artifact, and germline heterozygous in HG002 using trio sequencing data (Supplementary Fig. 2A). First, the HG002 300× Illumina BAM file and 250× Element BAM file were processed using prancSTR and BulkMonSTR’s mosaic genotyping. Candidate loci were initially selected based on the presence of at least three reads supporting the mutant allele in the orthogonal sequencing data; loci that did not meet this requirement were labeled as artifacts. Next, we required that the VAF of the mutant allele in the parental sequencing data be below 0.2. Loci with a VAF exceeding this threshold were classified as germline heterozygous. Furthermore, considering the prevalence of recurrent noise in repeat regions, we hypothesized that if the VAF of the mutant allele in the called samples was not significantly higher than the VAF observed in both parents’ sequencing data (binomial test with *P* > 0.05), the locus should be flagged as an artifact. Loci passing all these steps were designated as candidate mosaic STR mutations. To construct a high-confidence ground truth for HG002 mosaic STR mutations, all candidate loci were manually reviewed using IGV (v2.11.9). Ultimately, we labeled 129 indel mosaic mutations, 24 mismatch mosaic mutations, 51,539 indel germline heterozygous, 5,205 mismatch germline heterozygous, 107,960 indel artifacts, and 47,036 mismatch artifacts. These loci were then downsampled to various depths (30×, 50×, 100×, 150×, 200×, 250×, and 300×) to facilitate model training across different sequencing depths.

### Simulation of mosaic STR mutations

The workflow for simulating mosaic STR mutations is outlined in Supplementary Fig. 2B. To generate robust simulation data, STR loci meeting predefined criteria are selected for simulation (Supplementary Fig. 2B). These loci, along with mutation models, stutter error models, a reference genome FASTA, and the HG002 BAM file, serve as inputs for the simulation. Mutation types, sites, and lengths are sampled according to the mutation model. For indels, a stepwise mutation model is used^39,40^, incorporating insertion (0.5) and deletion (0.5) rates, as well as a geometric distribution (parameterized by *ρ*; homopolymer *ρ* = 0.8; other motif lengths *ρ* = 0.9) to control mutation size. Indel positions are selected to maintain the continuity of tandem repeats. For mismatches, a simplified K2P-like mutation model^41^ is applied, with transition/transversion rates set to 5:2 and mutation positions randomly chosen. Stutter errors are then introduced based on a stepwise error model, derived from our estimated per-locus stutter error model using population-level data. This enables the simulation of stutter error spectra for each allele. Finally, BamSurgeon^23^ (v1.4.1) introduces spike-in mosaic STR mutations, and the resulting BAM files are mixed to achieve the desired sequencing depths for all simulated alleles. This process yields two distinct simulation datasets (Supplementary Table S3):

1. SimData1 (Genotyping evaluation dataset / Training dataset): SimData1 is used for both genotyping evaluation and training the BulkMonSTR RF model. It consists of 20,766 loci for mismatch simulation (3,461 loci per motif length) and 38,400 loci for indel simulation (6,400 loci per motif length). VAFs are simulated at levels of 0.02, 0.03, 0.05, 0.1, and 0.3, following an allocation ratio of 4:4:4:2:1. Based on these VAF values, mutant allele depths are generated using binomial sampling. The simulated mutations are sampled to seven depths: 30×, 50×, 100×, 150×, 200×, 250×, and 300×. In total, SimData1 includes 114,885 mosaic STR mismatches and 206,896 mosaic STR indels.
2. SimData2 (Evaluation dataset): SimData2 is designed for model evaluation. It contains 6,000 STR loci (1,000 loci per motif length) for mismatch simulations, and an additional 6,000 STR loci (1,000 loci per motif length) for indel simulations. This dataset covers a broader range of VAFs (0, 0.01, 0.02, 0.03, 0.05, 0.1, 0.15, 0.2, 0.25, 0.3, 0.4, 0.5) and includes varying stutter error rates (0.01, 0.05, 0.1). It also encompasses the same depth range as SimData1 (30×, 50×, 100×, 150×, 200×, 250×, and 300×). Ultimately, SimData2 contains 151,200 mosaic STR mismatches and 151,200 mosaic STR indels.

### Population allele frequency annotation

We obtained a population genotyping panel for tandem repeat variation by downloading EnsembleTR calls^20,21^ on the 1000 Genomes Project and H3Africa cohorts (Version II). This panel, which comprises 3,550 individuals, integrates data from four different tools: HipSTR^19^, GangSTR^42^, adVNTR^43^, and ExpansionHunter^44^. Given that sequence-based genotyping is provided in the VCF files, we extracted all alleles for each STR locus within the population and calculated their frequencies, which were then stored in a BED file. This resource was subsequently used to annotate population allele frequencies of mosaic STR mutations and to filter potential germline variants.

### Filtering mapping artifacts by excluding segmental duplication STR loci

Segmental duplication (SD) can cause mapping artifacts, leading to a significant enrichment of mutation clusters within STR regions. These artifacts are difficult to distinguish from mosaic mutations. To address this, we excluded STR loci located within SD regions. However, after filtering SD regions based on the GIAB GRCh38 SD (v3.5) panel^45^, we still observed mutation clusters in some loci. We suspected that the SD regions in GRCh38 might be incomplete, as the latest T2T reference genome reports a higher proportion of SDs (7.0% compared to 5.4% in GRCh38)^46^. To test it, we extracted the reads with mutation clusters and re-aligned them to the T2T reference genome. We found that the mutation clusters were absent in the T2T alignments (Supplementary Fig. 14), confirming our hypothesis. Furthermore, we found that these mutation clusters did indeed occur within T2T-specific SD regions. Therefore, we downloaded T2T segmental duplication data^46^ from UCSC and performed a lift-over to the target reference genome version. This data serves as a supplement to the non-T2T SD regions from GIAB, adding 6,715 additional STR loci within SD regions for GRCh37 and 9,570 for GRCh38.

### Heuristic filtering of candidate STR mutations

To further refine BulkMonSTR calls, we applied heuristic filters to remove residual false positives. These filters operate downstream of the random forest classifier and apply permissive thresholds across read-support, mapping-level, sequencing-level, population-level, and stutter-related metrics. Filtering criteria and corresponding thresholds are provided in Supplementary Table S1.

### BulkMonSTR case-control mode

BulkMonSTR was adapted for somatic STR mutation calling in a paired case-control design. Case and matched control samples were processed jointly, with mosaic and germline genotypes estimated for each sample. A somatic STR mutation is identified when an allele is present in the case sample but absent from the control sample’s germline genotype. Loci classified as artifacts by the BulkMonSTR RF model were excluded. To exclude residual germline variants and recurrent technical noise, the following filters were applied: (1) Mutations with VAF ≥ 20% in the matched control sample were removed. (2) Mutations for which mutant allele read counts in the case sample were not significantly higher than those in the control sample were removed using an exact binomial test with *P* > 0.05. (3) Mutations for which mutant allele read counts in the case sample were not significantly higher than unassigned read counts were removed using an exact binomial test with *P* > 0.05.

### Detection of STR mutations using established algorithms

prancSTR^18^ (v6.0.1) was run to detect mosaic STR mutations without a matched normal. Following the recommended pipeline, we first ran HipSTR^19^ (v0.7) with default parameters, except for specifying --min-reads 10 to accommodate genotyping across various depths. We also used --stutter-in to input a stutter model estimated from HG002, ensuring consistency with BulkMonSTR. Next, dumpSTR^47^ (v6.0.1) was used to filter out regions of segmental duplications and low mappability from GIAB^45^ (keep consistency with BulkMonSTR) by specifying a bed file via the --filter-regions parameter. prancSTR was then executed with the recommended parameters: --vcftype hipstr, --readfield MALLREADS, and --only-passing. The output of prancSTR was filtered to include candidate mosaic STR mutations with the following criteria: at least three reads supporting the identified mosaic allele, a read depth of at least 10, 0.02 < VAF ≤ 0.3, HipSTR quality score ≥ 0.8, and Benjamini-Hochberg-corrected p-value (padj) < 0.05. For 170 TCGA blood samples, mosaic STR mutations detected in >10 samples were removed. Samples with mutation counts exceeding two standard deviations above the cohort mean were excluded.

The mutation list for HG008^34^ tumor samples was downloaded from the GIAB FTP site (https://ftp.ncbi.nlm.nih.gov/ReferenceSamples/giab/data_somatic/HG008/Liss_lab/analysis). The complete list of downloaded mutation files is available in Supplementary Table S2, and all mutation sites within STR regions are listed in Supplementary Table S6. We obtained mutation lists from short-read callers, including Mutect2^16^, Lancet^35^, and Strelka2^17^. The downloaded mutation list was a merged set, where high-confidence mutations were designated as those identified by at least two tools. Additionally, mutation lists from DeepSomatic^12^ and ClairS^14^ were downloaded and run on PacBio HiFi data. The workflow and parameters used to run these tools are provided in the corresponding README.md files on the GIAB FTP site. All mutations were then restricted to STR regions, with loci in segmental duplication and low mappability regions excluded. For control-independent benchmarking, analyses were further restricted to STR loci outside tumor copy number variation (CNV) regions to avoid simulation bias of tumor purities. For case–control analyses, genome-wide STR loci were retained. Tumor CNV annotations were obtained from the GIAB FTP site (https://ftp-trace.ncbi.nlm.nih.gov/ReferenceSamples/giab/data_somatic/HG008/Liss_lab/analysis/NIH_HiFi-HiC_Wakhan-CNA_20240424/bed_output).

### Construction of *in silico* mixed bulk sequencing data for HG008

High-depth Element sequencing data for HG008 were obtained from the GIAB Consortium FTP repository (Supplementary Table S2). Multiple datasets generated using standard insert libraries (350–400 bp) and long-insert libraries (>1 kb) were combined to produce aggregated datasets with effective depths of 331× for PDAC tissue and 237× for the matched normal duodenal tissue. To construct *in silico* mixed bulk sequencing data, tumor and normal datasets were combined at predefined proportions (5%, 10%, 20%, 40%, 80%, and 100%), and subsequently downsampled to a range of sequencing depths (30×, 50×, 100×, 150×, 200×, and 250×) for benchmarking analyses.

### Methodology for validation of candidate mosaic STR mutations

To validate candidate mosaic STR mutations, we employed two distinct methods. The first approach leverages paired sequencing data together with an orthogonal sequencing platform. (Supplementary Fig. 15A), while the second approach is based on read-based phasing (Supplementary Fig. 15B).

#### (1) Validation using paired and orthogonal sequencing data

Candidate STR mutations in HG005 and HG008 were validated using paired and orthogonal sequencing data (Supplementary Table S5). For HG005, the paired data refer to Illumina sequencing data from the father (HG006) and mother (HG007). For HG008, the paired data correspond to sequencing data from the matched normal tissue. Orthogonal sequencing data refer to data generated from the same sample using a different sequencing platform (i.e., Illumina data validated using Element sequencing, and Element data validated using Illumina sequencing). Compared to target sequencing, high-depth cross-platform WGS offers higher throughput for validating genome-wide STR loci. The inclusion of paired sequencing data was crucial for effectively labeling false positives from germline heterozygous variants and recurrent artifacts. The validation steps were as follows: (i) Candidate loci were required to have at least two reads supporting the mutant allele in the orthogonal Element/Illumina sequencing data. Loci failing this criterion were labeled as artifacts. (ii) The variant allele frequency of the mutant allele in the paired sequencing data was required to be below 0.2. Loci with a paired VAF above this threshold were classified as germline heterozygous variants. (iii) Given the prevalence of recurrent noise in repeat regions, we hypothesized that if the mutant allele’s VAF in the called sample was not significantly higher than the VAF of either parent’s sequencing data (binomial test with Benjamini correction, adjusted p > 0.05), the locus would be considered a recurrent artifact. Only those loci that successfully passed all of these validation steps were deemed true positive mosaic STR mutations.

#### (2) Validation using read-based phasing

The TCGA blood samples (all sample information is available in Supplementary Table S7) were validated using read-based phasing (Supplementary Table S8). While it is restricted to phasable loci (specifically those with a germline heterozygous SNP near the STR loci), we considered it an unbiased approach for comparing the performance of different tools. Given the assumption that mosaic mutations are rare, it is highly unlikely that they occur independently at the same genomic position on both the maternal and paternal copies of the genome. Therefore, each phasable candidate mosaic mutation was classified as follows: (i) Loci that resulted in three haplotypes were classified as mosaics (hap=3). (ii) Loci that resulted in two haplotypes were classified as germline heterozygous variants (hap=2). (iii) Loci that resulted in more than three haplotypes were classified as artifacts (hap>4). To perform read-based phasing, we obtain germline heterozygous SNPs of TCGA blood samples. The BAM files were processed using the GATK (v4.2.4.1)^48^ best practice workflow. Strict filtering criteria were applied to ensure high-quality germline SNPs, including: (i) excluding VQSR FAIL variants, (ii) using a two-tailed binomial test to remove variants deviating from a VAF of 0.5, (iii) filtering out variants with genotype quality ≤20, and (iv) excluding variants with a depth <20. To reduce the impact of random events on the accuracy of haplotype counting, only high-quality phasable loci were used, which met the following criteria: (i) at least 10 phasable reads, (ii) at least two reads supporting each phased haplotype. After validation using these methods, all confirmed mutations from different tools were compiled into a sample-specific ground truth set, which was subsequently used to assess the performance of each tool.

### Profile recurrent errors within STR regions

To characterize the recurrent error profile of STR regions, we used HG002, a widely adopted benchmark sample with a highly accurate genome assembly^49^. By leveraging this well-characterized sample, we could precisely quantify and characterize the types of noise that commonly occur in STR regions. Given the high noise levels in STR regions and the rarity of mosaic mutations, we hypothesized that the most recurrent non-germline alleles, supported by at least two reads, are likely artifacts that could significantly affect mosaic mutation calling. To capture these recurrent artifacts, we focused on loci with unambiguous germline alleles, applying the following stringent criteria: (1) homozygous sites, to minimize the impact of confounding unknown noise sources; (2) loci located within well-assembled genomic regions, where STR allele sequences are confidently assembled; and (3) exclusion of loci within segmental duplication regions to avoid mapping errors due to the limitations of short-read sequencing. Once these recurrent artifact alleles were identified from the specified loci, we inferred that their origin was the known germline allele. By comparing these artifacts to the corresponding germline allele, we classified them into four types: SNVs, MNVs, insertions, and deletions. We then characterized the error profiles and features of these identified artifacts for both Illumina and Element sequencing platforms.

### Estimation of the stutter model from different sequencing platforms in HG002

To estimate the stutter error of the GIAB HG002 sample, we first assumed that loci with the same motif length and similar total length exhibit similar stutter error rates, and that mutations are rare across these loci. Next, homozygous STR loci with a determined sequence in the assembled regions were selected. Sequencing reads with an allele length different from the assembled sequence were considered stutter errors. Based on this, a generalized stepwise error model (Equation 1) was used to construct a stutter model for different motif lengths within specific STR total length ranges. The insertion rate corresponds to the proportion of all stutter insertion reads, the deletion rate represents the proportion of all stutter deletion reads, and the geometric step-size distribution parameter reflects the proportion of stutter reads relative to the total indel length change. Stutter errors that are integer multiples of the motif length are defined as in-frame stutter errors, while non-integer multiples are defined as out-of-frame stutter errors. Using this approach, in-frame and out-of-frame stutter error models were constructed for different sequencing technologies. A simplified, motif-length-based stutter error model was then used to visualize the stutter errors across different sequencing technologies (Supplementary Fig. 16, Supplementary Table S11).

### Mutation signature analysis

The mutational signatures of mosaic STR mutations in 170 TCGA blood samples were analyzed. Mutation matrices were generated using SigProfilerMatrixGenerator^50^ (v1.2.31), and known COSMIC mutational signatures were subsequently assigned to each sample using SigProfilerAssignment^29^ (v0.1.9) based on a forward stagewise algorithm and non-negative least squares.

### Functional annotation of STR mutations

ANNOVAR^51^ (version 2020-06-07) was used to annotate exonic variants, associated genes, functional impact, and predicted protein-coding consequences. Tissue-specific cis-regulatory element annotations from ENCODE (version 4, hg38)^52^ were downloaded from the SCREEN (Search Candidate Regulatory Elements by ENCODE) platform (https://screen.wenglab.org/downloads). STR mutation sites identified in HG002, HG005, and TCGA blood samples were annotated using candidate cis-regulatory elements (cCREs) derived from blood tissues. STR mutations in HG008 were annotated using pancreas-specific cCRE annotations (Supplementary Table S4, Supplementary Table S8-10).

### Data access

Information on all sequencing datasets and publicly available resources used in this study, including their download links, is provided in Supplementary Table S2. Simulated mosaic STR mutations are listed in Supplementary Table S3. Validated STR mutations and corresponding validation results of GIAB samples are provided in Supplementary Tables S4-S5. HG008 mutation calls from established small variant callers were downloaded from the GIAB FTP resource and are summarized in Supplementary Table S6. A summary of TCGA blood samples included in this study is provided in Supplementary Table S7, with corresponding STR mutation lists available in Supplementary Table S8. STR mutations within coding sequences are provided in Supplementary Table S9. Comprehensive STR mutation analyses of all samples are summarized in Supplementary Table S10. Parameters of the HG002 stutter model estimated across sequencing platforms are reported in Supplementary Table S11. Additional resources for feature computation and mutation filtering include segmental duplication regions (with T2T SD panel lift-over intervals), processed EnsemblTR calls with population allele frequencies from 3,550 individuals, and STR locus mappability scores, all of which are available at GitHub (https://github.com/douymLab/BayesMonSTR/tree/main/BayesMonSTR-BulkMonSTR).

## Software availability

BulkMonSTR is written in Python and is licensed under the MIT License. The source code, documentation, examples, and conda configuration file are available on GitHub at https://github.com/douymLab/BayesMonSTR/tree/main/BayesMonSTR-BulkMonSTR. The source code of BulkMonSTR is also available as Supplemental Code.

## Acknowledgments

We would like to thank the GIAB community for constant support. This work was supported by the Westlake Laboratory of Life Sciences and Biomedicine (Hangzhou 310024, Zhejiang, China) under the grant “Key R&D Program of Zhejiang” (2024SSYS0032) as well as the National Natural Science Foundation of China (32270682) to Y.D.. We thank the High-Performance Computing Center for their technical support. We also acknowledge financial support from the National Natural Science Foundation of China (grant number: 32270682) and the Westlake Education Foundation. We would like to thank the submitters of the following datasets: GIAB reference sample sequencing data from the GIAB Consortium, TCGA blood samples under dbGaP accession phs000178.v11.p8, and Element sequencing data from Google Brain. The public datasets were obtained from the following sources: GIAB at https://www.nist.gov/programs-projects/genome-bottle, dbGaP at http://www.ncbi.nlm.nih.gov/gap, and Google Brain at https://console.cloud.google.com/storage/browser/brain-genomics-public/research. Certain commercial equipment, instruments or materials are identified to adequately specify the experimental conditions or reported results. Such identification does not imply recommendation or endorsement by the National Institute of Standards and Technology, nor does it imply that the equipment, instruments or materials identified are necessarily the best available for the purpose.

## Author contributions

Y.D. conceived and supervised the project and secured the funding. W.W. developed the BulkMonSTR and performed the analysis with help from W.L., C.W., W.F. and Y.X.. X.Y. and C.C. contributed to data access and discussion. W.W. and Y.D. wrote the manuscript.

## References

1. Levinson, G. & Gutman, G. A. Slipped-strand mispairing: a major mechanism for DNA sequence evolution. Mol. Biol. Evol. 4, 203–221 (1987).

2. Porubsky, D. et al. Human de novo mutation rates from a four-generation pedigree reference. Nature 1–10 (2025) doi:10.1038/s41586-025-08922-2.

3. Fotsing, S. F. et al. The impact of short tandem repeat variation on gene expression. Nat Genet 51, 1652–1659 (2019).

4. Horton, C. A. et al. Short tandem repeats bind transcription factors to tune eukaryotic gene expression. Science 381, eadd1250 (2023).

5. Tanudisastro, H. A. et al. Polymorphic tandem repeats shape single-cell gene expression across the immune landscape. bioRxiv 2024.11.02.621562 (2025) doi:10.1101/2024.11.02.621562.

6. Manigbas, C. A. et al. A phenome-wide association study of tandem repeat variation in 168,554 individuals from the UK Biobank. Nat. Commun. 15, 10521 (2024).

7. Margoliash, J. et al. Polymorphic short tandem repeats make widespread contributions to blood and serum traits. Cell Genom. 3, 100458 (2023).

8. Rajan-Babu, I.-S., Dolzhenko, E., Eberle, M. A. & Friedman, J. M. Sequence composition changes in short tandem repeats: heterogeneity, detection, mechanisms and clinical implications. Nat. Rev. Genet. 1–24 (2024) doi:10.1038/s41576-024-00696-z.

9. Erwin, G. S. et al. Recurrent repeat expansions in human cancer genomes. Nature 1–7 (2022) doi:10.1038/s41586-022-05515-1.

10. Raz, O. et al. Short tandem repeat stutter model inferred from direct measurement of in vitro stutter noise. Nucleic Acids Res. 47, 2436–2445 (2019).

11. Stoler, N. & Nekrutenko, A. Sequencing error profiles of Illumina sequencing instruments. Nar Genom Bioinform 3, lqab019– (2021).

12. Park, J. et al. Accurate somatic small variant discovery for multiple sequencing technologies with DeepSomatic. Nat. Biotechnol. 1–10 (2025) doi:10.1038/s41587-025-02839-x.

13. Yang, X. et al. Control-independent mosaic single nucleotide variant detection with DeepMosaic. Nat Biotechnol 1–8 (2023) doi:10.1038/s41587-022-01559-w.

14. Zheng, Z. et al. ClairS: a deep-learning method for long-1read somatic small variant calling. bioRxiv (2023) doi:10.1101/2023.08.17.553778.

15. Dou, Y. et al. Accurate detection of mosaic variants in sequencing data without matched controls. Nat Biotechnol 38, 314–319 (2020).

16. Cibulskis, K. et al. Sensitive detection of somatic point mutations in impure and heterogeneous cancer samples. Nat Biotechnol 31, 213–219 (2013).

17. Kim, S. et al. Strelka2: fast and accurate calling of germline and somatic variants. Nat Methods 15, 591–594 (2018).

18. Sehgal, A., Jam, H. Z., Shen, A. & Gymrek, M. Genome-wide detection of somatic mosaicism at short tandem repeats. Bioinformatics 40, btae485 (2024).

19. Willems, T. et al. Genome-wide profiling of heritable and de novo STR variations. Nat Methods 14, 590–592 (2017).

20. Jam, H. Z. et al. A deep population reference panel of tandem repeat variation. Nat. Commun. 14, 6711 (2023).

21. Lundström, O. S. et al. WebSTR: a population-wide database of short tandem repeat variation in humans. Journal of Molecular Biology 168260 (2023) doi:10.1016/j.jmb.2023.168260.

22. Zook, J. M. et al. Extensive sequencing of seven human genomes to characterize benchmark reference materials. Sci Data 3, 160025 (2016).

23. Ewing, A. D. et al. Combining tumor genome simulation with crowdsourcing to benchmark somatic single-nucleotide-variant detection. Nat Methods 12, 623–630 (2015).

24. Daniels, C. A. et al. Characterization of subclonal variants in HG002 Genome in a Bottle reference material as a resource for benchmarking variant callers. Cell Genom. 101104 (2025) doi:10.1016/j.xgen.2025.101104.

25. Muzny, D. M. et al. Comprehensive molecular characterization of human colon and rectal cancer. Nature 487, 330–337 (2012).

26. Bohrson, C. L. et al. Linked-read analysis identifies mutations in single-cell DNA-sequencing data. Nat Genet 51, 749–754 (2019).

27. Ju, Y. S. et al. Somatic mutations reveal asymmetric cellular dynamics in the early human embryo. Nature 543, 714–718 (2017).

28. Lundberg, S. & Lee, S.-I. A Unified Approach to Interpreting Model Predictions. arXiv (2017) doi:10.48550/arxiv.1705.07874.

29. Díaz-Gay, M. et al. Assigning mutational signatures to individual samples and individual somatic mutations with SigProfilerAssignment. Bioinformatics 39, btad756 (2023).

30. Alexandrov, L. B. et al. The repertoire of mutational signatures in human cancer. Nature 578, 94–101 (2020).

31. Carroll, A. et al. Accurate human genome analysis with element avidity sequencing. BMC Bioinform. 26, 194 (2025).

32. Happ, H. C., Sasani, T. A., Warner, D., Neklason, D. W. & Quinlan, A. R. AVITI sequencing of a four-generation CEPH/Utah pedigree confirms low mutation rates at homopolymer loci despite their low sequence complexity. bioRxiv 2025.09.25.678675 (2025) doi:10.1101/2025.09.25.678675.

33. Zhang, X. et al. Systematic evaluation of the impact of promoter proximal short tandem repeats on expression. (2025) doi:10.1101/2025.09.14.676153.

34. McDaniel, J. H. et al. Development and extensive sequencing of a broadly-consented Genome in a Bottle matched tumor-normal pair. Sci. Data 12, 1195 (2025).

35. Narzisi, G. et al. Genome-wide somatic variant calling using localized colored de Bruijn graphs. Commun Biology 1, 20 (2018).

36. Genome in a Bottle Consortium. Somatic SNV and InDel callsets for HG008 generated by Mutect2, Strelka2, Lancet, ClairS, and DeepSomatic. https://ftp-trace.ncbi.nlm.nih.gov/ReferenceSamples/giab/data_somatic/HG008/Liss_lab/analysis.

37. Li, H. & Durbin, R. Fast and accurate short read alignment with Burrows–Wheeler transform. Bioinformatics 25, 1754–1760 (2009).

38. Wang, C. et al. Unravelling genome-wide mosaic microsatellite mutations at single-cell resolution. bioRxiv 2026.02.04.703915 (2026) doi:10.64898/2026.02.04.703915.

39. Willems, T. et al. Population-Scale Sequencing Data Enable Precise Estimates of Y-STR Mutation Rates. Am J Hum Genetics 98, 919–933 (2016).

40. Gymrek, M., Willems, T., Reich, D. & Erlich, Y. Interpreting short tandem repeat variations in humans using mutational constraint. Nat Genet 49, 1495–1501 (2017).

41. Kimura, M. A simple method for estimating evolutionary rates of base substitutions through comparative studies of nucleotide sequences. J. Mol. Evol. 16, 111–120 (1980).

42. Mousavi, N., Shleizer-Burko, S., Yanicky, R. & Gymrek, M. Profiling the genome-wide landscape of tandem repeat expansions. Nucleic Acids Res 47, e90–e90 (2019).

43. Park, J., Bakhtiari, M., Popp, B., Wiesener, M. & Bafna, V. Detecting tandem repeat variants in coding regions using code-adVNTR. Iscience 25, 104785 (2022).

44. Dolzhenko, E. et al. Detection of long repeat expansions from PCR-free whole-genome sequence data. Genome Res 27, 1895–1903 (2017).

45. Dwarshuis, N. et al. The GIAB genomic stratifications resource for human reference genomes. Nat. Commun. 15, 9029 (2024).

46. Vollger, M. R. et al. Segmental duplications and their variation in a complete human genome. Science 376, eabj6965 (2022).

47. Mousavi, N. et al. TRTools: a toolkit for genome-wide analysis of tandem repeats. Bioinformatics 37, 731–733 (2020).

48. DePristo, M. A. et al. A framework for variation discovery and genotyping using next-generation DNA sequencing data. Nat Genet 43, 491–498 (2011).

49. Hansen, N. F. et al. A complete diploid human genome benchmark for personalized genomics. bioRxiv 2025.09.21.677443 (2025) doi:10.1101/2025.09.21.677443.

50. Bergstrom, E. N. et al. SigProfilerMatrixGenerator: a tool for visualizing and exploring patterns of small mutational events. BMC Genom. 20, 685 (2019).

51. Wang, K., Li, M. & Hakonarson, H. ANNOVAR: functional annotation of genetic variants from high-throughput sequencing data. Nucleic Acids Res. 38, e164–e164 (2010).

52. Moore, J. E. et al. An expanded registry of candidate cis-regulatory elements. Nature 1–10 (2026) doi:10.1038/s41586-025-09909-9.

